# CD300f immune receptor is a microglial tissue damage sensor and regulates efferocytosis after brain damage

**DOI:** 10.1101/2024.09.06.611174

**Authors:** María Luciana Negro-Demontel, Frances Evans, Andrés Cawen, Zach Fitzpatrick, Hannah D. Mason, Daniela Alí, Hugo Peluffo

## Abstract

Microglia, the resident phagocytes of the central nervous system (CNS), continuously monitor the parenchyma and surrounding borders and are the primary responders to brain damage. CD300f is a lipid-sensing immunoreceptor present in the microglial cell membrane, which binds to phosphatidylserine and other lipid mediators. Defining the functional microglial sensome is critical to understand their function and cell state determination. Using intravital two-photon microscopy we show that microglia lacking the CD300f receptor fail to detect environmental damage cues after a laser ablation injury. After a mild traumatic brain injury or after the intracortical injection of apoptotic cells, CD300f^-/-^ microglia showed reduced capacity for detecting and phagocytosing dyeing cells, leading to the accumulation of dead cells in the neural parenchyma. Moreover, at later timepoints, increased accumulation of dyeing cells was found inside CD300f^-/-^ microglia in vivo and in bone marrow-derived macrophages in vitro, suggesting that these cells display a reduced capacity for metabolizing phagocytosed cells. Finally, CD300f deficiency increased functional compromise after a contusive traumatic brain injury, associated to increased conservation of brain tissue. Collectively, these results suggest that CD300f function as a damage-associated molecular pattern (DAMP) receptor that coordinates microglial process reaction towards tissue debris and highlights its central role in microglial sensome machinery and in the modulation of in vivo microglial efferocytosis.

## Introduction

Microglia are innate immune cells that reside in the central nervous system (CNS) parenchyma. Pioneering work from 15 years ago demonstrated that microglia actively sense the neural parenchyma ^1,2^. Subsequent descriptions of the microglial transcriptome and its sensome ^3^ have contributed to the identification and understanding of essential functions of microglia related to their sensing functions, such as synapse pruning, detection of tissue damage and pathogens, and engulfment of apoptotic cells, myelin debris, and toxic lipids and proteins such as Aβ ^4^.

Phagocytosis is, in many cases, an essential step that follows microglial sensing of damage-associated molecular patterns (DAMP) or apoptotic cells, and microglial purine receptors ^1^ as well as TAM receptors ^5^serve as central mediators of these processes. Moreover, microglia have been shown to express lipid-sensing immune receptors such as TREM2, which directly participate in the recognition and phagocytosis of apoptotic cells and tissue debris such as myelin ^6,7^. It has been proposed that TREM2 could additionally act as a DAMP receptor or pathogen-associated molecular pattern (PAMP) receptor due to its ability to recognize phospholipids such as phosphatidylserine or sphingomyelin, lipoproteins, LPS, and Aβ (5). The CD300 family of lipid-binding immune receptors is less understood, but their members share most of their known ligands and signaling pathways with TREM2. CD300f is a unique receptor that integrates activating and inhibitory signals, and several lines of evidence suggest that it could act as a DAMP sensor. CD300f directly associates with CD300b negatively regulating its activity ^8^, and CD300b has been shown to engage LPS and, as an activating receptor, contribute to septic shock ^9^. Moreover, CD300f directly inhibits Toll-like receptors 2, 3, 4 and 9 signaling ^10^, and has been reported to interact with murine norovirus ^11^, showing that CD300f modulate diverse PAMP receptors or could act as a PAMP receptor itself. As CD300f is essential for immunometabolic reprogramming of microglia and macrophages ^12,13^, it opens the question whether this receptor would play a role in the microglial sensome by coordinating diverse components of the neuroimmune response and immunometabolism.

Large amounts of DAMPs are produced after traumatic CNS injuries. A contusion to the spinal cord leads to the induction of a specific microglia population that is not observed in degenerative disease, termed injury-activated microglia and macrophages (IAM) ^14^. Using INTACT RNA-seq and scRNA seq technology they revealed that there are 14 injury response genes, including *cd300f*, that intersected with IAM core signature genes that were uniformly expressed in IAM and in all reactive microglia and macrophage subclusters ^14^. These findings highlight that CD300f might be indispensable for modulating the responses of all microglia and macrophages population in the injured SCI.

CD300f has been shown to participate in synapse pruning in vitro by a phosphatidylserine (PS)-dependent mechanism ^12,15^ in a similar way as described for TREM2 ^16^. In accordance, knockout mice for CD300f exhibit sex-dependent depressive-like ^12^ and anxiety-like behaviors ^17^, and a single nucleotide polymorphism (SNP) for CD300f protects women from major depressive disorder ^12^ and men from generalized anxiety disorder ^17^. CD300f is also important for macrophage PS-mediated engulfment of apoptotic cells in vitro ^18^. In the brain, CD300f^-/-^ microglia show reduced levels of SPP1 ^12^, a key mediator of phagocytosis, suggesting that these cells may exhibit altered clearance capacities in the CNS parenchyma. In addition, CD300f not only physically interacts with IL4Rα, but is necessary for robust IL-4 responses ^19^ which are critical signals for macrophages and their contributions to tissue repair and fibrosis ^20^. Sensing of apoptotic cells, combined with IL-4 signaling, launches an anti-inflammatory and tissue repair response in macrophages in a wide range of biological contexts ^21^. Such work highlights a pivotal role for CD300f in various phagocytic mechanisms that extend beyond PS detection.

CD300f is emerging as an important immune receptor regulating microglia and macrophage function in many physiological and pathological conditions. For instance, CD300f knockout mice show accelerated inflammaging, cognitive decline, brain senescence and frailty, suggesting that it is important for healthy aging ^13^. These mice also show enhanced neurological impairment in the experimental autoimmune encephalomyelitis (EAE) multiple sclerosis model ^22^, and some SNPs have been associated to the disease ^23,24^. Its overexpression was neuroprotective after an acute brain injury ^25^, and has been correlated with improved neurological performance in Tau and Aβ AD models ^26,27^. However, the underlying mechanisms that dictate the roles CD300f plays in microglial responses to brain injury or under physiological conditions remain poorly understood. We report that CD300f is essential for the prompt response of microglial cells towards tissue damage and contributes to the detection, engulfment and processing of apoptotic cells in vivo. Under conditions where increased apoptotic cells and DAMPs are generated, such as in traumatic brain injury (TBI), microglial CD300f deficiency results in compromised functional recovery.

## Results

Defining the microglial sensome would be a critical step toward understanding the functions of these cells in the CNS. Lipid-binding immune receptors could function as sensors of CNS damage, coupling cell immunometabolism and general response to the changes in the extracellular milieu. We hypothesized that CD300f could constitute an important hub for recognizing extracellular damage signals and coordinating microglial phenotype reorganization. To explore this hypothesis, we generated CX3CR1^+/GFP^/CD300f^-/-^ mice, induced local DAMP production by a restricted brain laser lesion and followed the microglial reaction towards the lesion by two-photon microscopy. A significant reduction in microglial projection recruitment towards the lesion was observed in the absence of CD300f (Fig. 1A-C) (VIDEO 1), though important variations were observed between animals (Suppl. Fig. 1A). Interestingly, CD300f deficiency did not affect the basal dynamics of microglial processes in the absence of cell debris (Fig. 1D-E) nor their morphology (Suppl. Fig. 1B-D) (VIDEO 2). To further explore whether CD300f participates in microglial sensing of tissue damage and apoptotic cells, we performed a mild TBI (mTBI) to the cortex. Such an insult induces the morphological transformation of microglia towards phagocytic/jellyfish stages due to the local production of cell debris, purines, pyrimidines, and apoptotic cells, as described previously ^28^. After the induction of a mTBI in both WT and CD300f^-/-^ mice, dead cells were labelled by transcranial application of propidium iodide (PI), which permeate cell membranes of dead and dying cells within the lesion. We then quantified the shortest distances between microglial projections and a PI+ dying cells in the perilesion area of CD300f^-/-^ and WT brains (Fig. 1F-G). A significant increase in this distance was observed in the absence of CD300f, suggesting a reduced capability of CD300f^-/-^ microglia to sense and respond to dying cells. In accordance, when microglial cells under these conditions were followed by intravital two-photon microscopy, CD300f deficiency resulted in a reduction in phagocytic morphology (Fig. 1H-I) (VIDEO 3). Taken together, these results indicate that while the basal microglial surveillance of brain parenchyma remains unaffected in CD300f^-/-^ microglia, the absence of CD300f diminishes their sensitivity to local tissue damage cues and apoptotic cells.

**Figure 1.**
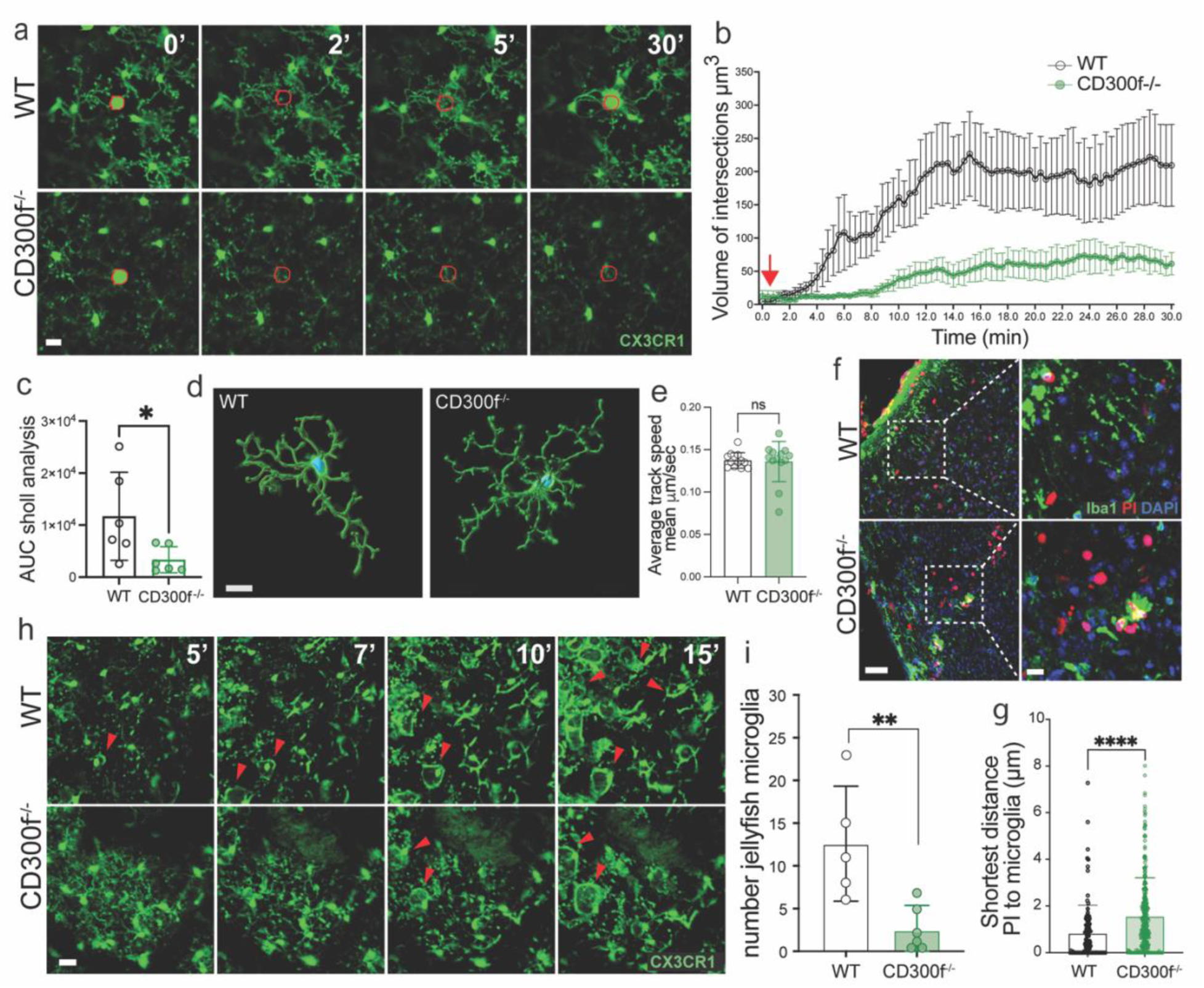
CD300f contributes to the microglial sensome. Representative two-photon timelapse stills through a thinned-skull window depict the acute response of microglia (CX3CR1^+/GFP^, shown in green) to laser-induced ablation. The site of laser impact is highlighted in red. Scale bar 10 μm **b.** Scholl analysis to measure microglial processes migration. WT microglial processes rapidly migrate toward the site of injury, whereas CD300f^-/-^ microglia fail to respond. Red arrow indicates the time where laser ablation was induced. Two-way ANOVA comparing animals, *p<0.05 WT (n=6, 10 lesions) vs CD300f^-/-^ (n=6, 12 lesions). **c**. The area under the curve (AUC) from plot b shows the overall magnitude of the microglial response following a laser-induced injury observed during the entire time course. Two-tailed Student’s t-test * p<0.05 **d.** Representative rendering of homeostatic microglia from intravital two-photon stacks, utilized for quantifying the average track speed of microglial processes. Scale bar 15 μm. Data presented in (**e**) demonstrates no alterations in microglial surveillance speed in the absence of the CD300f receptor. **f, g**. Confocal images of cortical brain showing an increase in the shortest distance of any given PI+ cell (red) to a microglial cell (green) in the perilesion area 5h post mTBI, n=5 (DAPI blue). Scale bars 40 μm and 10 μm. Two-tailed Student’s t-test ****p<0.0001. **h, i.** Intravital two-photon timelapse stacks of CX3CR1^+/GFP^ microglia illustrates morphological transformations and the induction of a phagocytic (jellyfish) phenotype minutes after mTBI in WT microglia. In contrast, CD300f-deficient microglia fail to induce this phagocytic morphology (arrowheads). Scale bar 20 μm. Two-tailed Student’s t-test**p<0.01 n=5 (WT) and n=6 (CD300f^-/-^).

It has been shown that macrophage CD300f is important for the phagocytosis of apoptotic cells in vitro ^18^. To explore whether microglial CD300f is involved in the efferocytosis of locally generated apoptotic cells in vivo, we analyzed the number of PI+ dead/dying cells accumulated inside CX3CR1+ microglia after a mTBI in both WT and CD300f^-/-^ mice (Fig. 2B-C). Intravital two-photon microscopy at 3 hours post mTBI demonstrated a reduction in the microglial phagocytosis rate in CX3CR1^+/GFP^/CD300f^-/-^ mice compared to WT mice (Fig. 2C). In these experiments, microglia dynamics were tracked for 30 minutes, starting at 3 hours after mTBI. CD300f deficient microglia exhibited an overall reduced phagocytic capacity evidenced by a decrease in intracellular PI+ signal at both the beginning and the end of the experiment, indicative of fewer actively engulfed dead cells (VIDEO 4). These findings are supported by histological analysis of CD300f^-/-^ brains after mTBI, demonstrating an increased accumulation of total PI+ cells in the center of the lesion at 5 hours post-injury (Fig. 2D-F).

**Figure 2.**
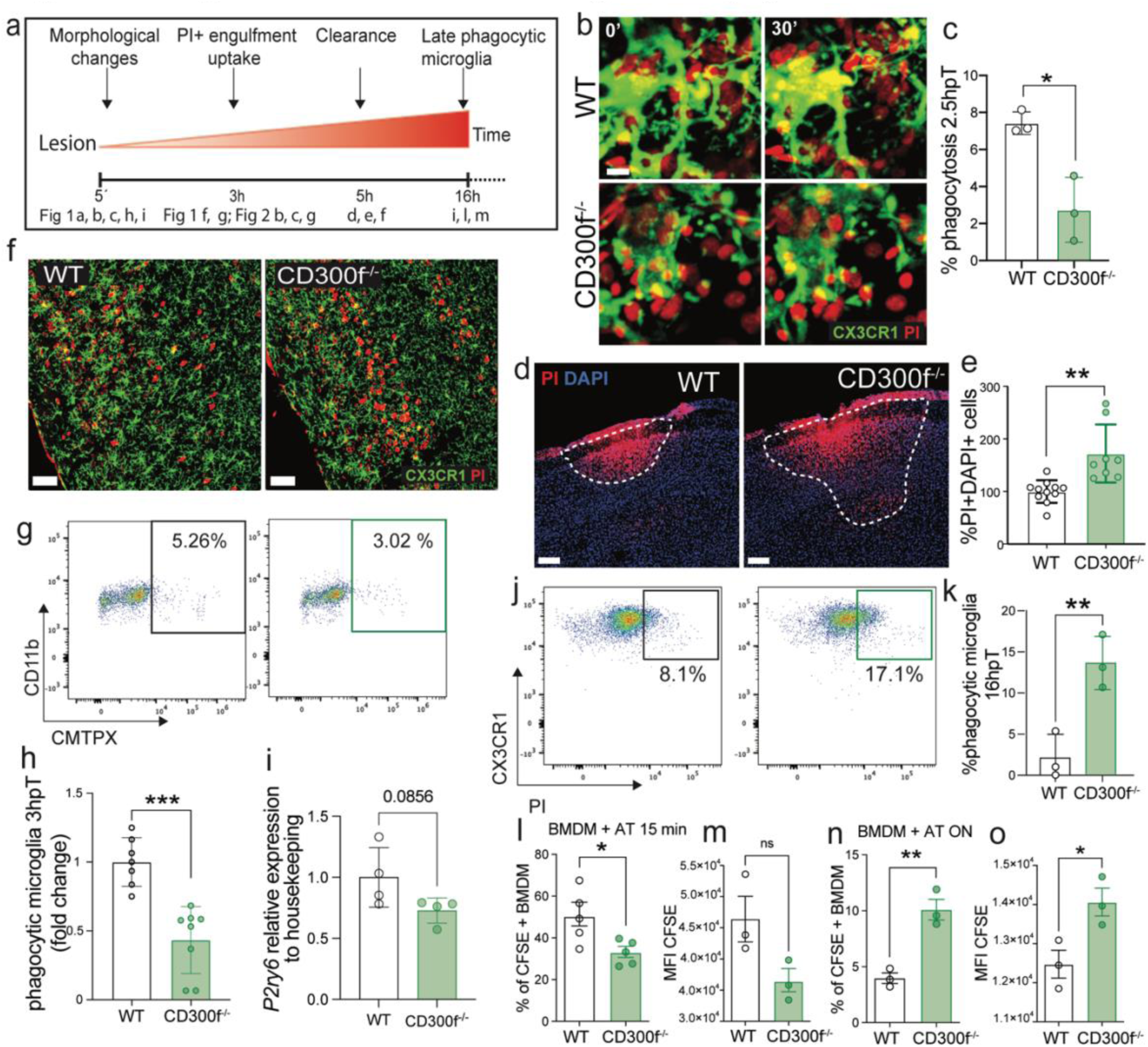
Microglia utilize CD300f receptor to detect and engulf dead/dying cells. **a.** Workflow schematic. **b, c**. Representative intravital two-photon microscopy stacks through a thinned-skull window show CX3CR1^+/GFP^ microglia engulfing dead cells (PI^+^, red) 3 hours after mTBI. The uptake rate of PI^+^ dead cells within a 30-minute timeframe by CD300f-deficient microglia is reduced after mTBI. Scale bar 10 μm. Statistical analysis was performed using a two-tailed Student’s t-test, *p<0.05, based on data pooled from 3 independent experiments, with one mouse per experiment (n=3 mice per group) **d.** Representative images depict CD300f^-/-^ and WT brains 5 hours after mTBI. Mice were treated with transcranial PI for 45 minutes prior to perfusion to label dead/dying cells. Scale bar 150 μm. Analysis in panel **e** shows an increase in PI^+^ dead/dying cell count in the brain parenchyma of CD300f-deficient mice compared to WT. Two-tailed Student’s t-test, **p<0.01, n=10 (WT), n=8 (CD300f^-/-^). **f.** Representative images showing WT vs CD300f^-/-^ microglia (green) in contact with PI+ dead/dying cells (red). Scale bar 50 μm. **g.** Reduced microglial phagocytosis of stained apoptotic thymocytes (AT) injected into the brain parenchyma of CD300f^-/-^ mice compared to WT animals, analyzed by flow cytometry of the lesioned brain area 5 hours after injection. Two-tailed Student’s t-test was conducted on data pooled from 3 independent experiments with n=7 (WT) and n=8 (CD300f^-/-^) mice per group. **h**. QPCR *P2ry6* relative expression in naïve WT vs CD300f^-/-^ brains. **i**. Flow cytometry of 2 mm^3^ brain punches was performed 16 hours post mTBI-PI treatment, and phagocytic microglia (gated on viable/CD45^+^/CD11b^low^/PI^+^) were analyzed. The graph shows an increase in CD300f^-/-^ late phagocytic microglia compared with WT, indicating a reduction/delay in dead cell clearance and/or an increase in rate of digestion. Data were normalized to phagocytic WT microglia (based on 3 independent experiments with 3-5 brain punches pooled together). **j-m**. CD300f^-/-^ and WT BMDMs were incubated for 15 minutes (**j, k**) or 4 hours (**l, m**) with stained AT. Short-term incubation of CD300f^-/-^ BMDMs with AT (**j, k**) demonstrates a decrease in uptake of AT, and geometric mean fluorescence intensity [gMFI] of AT in CD300f^-/-^ BMDM, while long-term incubation (**l, m**) reveals an accumulation of dead cells in CD300f^-/-^ BMDM cytoplasm (**l**, % of AT+BMDM and **m**, gMFI of AT in BMDM). Data are from 3 independent experiments. Two-tailed Student’s t-test, *p<0.05, **p<0.01. n=3 animals per group.

To further elucidate the in vivo role of CD300f in dead cell detection and clearance by microglial cells, we injected fluorescently labelled apoptotic thymocytes into the cortex of WT and CX3CR1^+/GFP^/CD300f^-/-^ mice as described previously ^29^ (Fig. 2G) (VIDEO 5). Quantification of microglial uptake of apoptotic thymocytes was performed after isolating cells from the cortex at 3 hours post-injection and processing them for flow cytometry analysis. A significant reduction in CD11b+ microglia containing engulfed apoptotic thymocytes was detected in the absence of CD300f. Purines are released during periods of damage, inflammation, and hyperexcitability, evoking calcium signaling in microglia predominantly through P2RY6 ^30^. Given that this receptor is restricted to microglia and hypothalamic neurons ^31^, and is key for microglial phagocytosis ^30,31^ and morphological switch to jellyfish/phagocytic conformation ^28^, we examined microglial levels of p2ry6 mRNA expression by QPCR of naïve cortex samples. We observed a strong tendency towards reduced expression in CD300f-/-brains (Fig. 2H), suggesting a potential contribution to the decreased sensing and phagocytotic capacity of CD300f^-/-^ microglia.

While a diminished capacity to sense and phagocytose dying cells was noted during the early time points after lesion formation, at 16 hours post-injury, an increase in the content of PI+ dying cells was observed in CD300f^-/-^ microglial cells (Fig. 2I). This could be explained by a delayed increase in efferocytosis of CD300f^-/-^ microglia, or alternatively, these cells could be unable to digest the phagocytosed material, in a similar manner as observed with TREM2-deficient microglia ^32^. To gain a deeper understanding of this phenomenon, we cultured bone marrow-derived macrophages (BMDM) from WT and CD300f^-/-^ mice and exposed them to labelled apoptotic thymocytes in vitro. We then assessed the efficiency of efferocytosis and the accumulation of apoptotic thymocytes within the macrophages. Consistent with the in vivo results, CD300f^-/-^ macrophages demonstrated reduced efferocytosis during short incubation periods with apoptotic thymocytes (Fig. 2J-K). However, CD300f-deficient macrophages displayed increased accumulation of phagocytosed thymocytes at later timepoints, suggesting a reduced capacity to degrade phagocytosed cells (Fig. 2L, M). Collectively, these findings suggest that CD300f deficient microglial cells not only fail to sense molecules derived from apoptotic cells, thus impairing their ability to engage with them effectively, but also possess an impaired capacity for processing engulfed cellular material.

The integrity of the glial limitans is pivotal in maintaining compartmentalized brain responses, as a breach could lead to an influx of danger signals from the periphery, potentially resulting in increased cell death. To evaluate the functional integrity of the glia limitans superficialis in WT and CD300f^-/-^ mice, we administered a low-molecular-weight fluorescent dye (SR101) transcranially. Typically, SR101 remains localized primarily to the meninges when the glia limitans superficialis is intact ^28^. However, a mTBI-induced breach in this structure allows SR101 passage into the neocortex, as assessed using intravital two-photon microscopy ^28,33^. Following mTBI, we observed no differences in glial limitans permeability between CD300f^-/-^ and WT brains (Suppl. 2A-B). This suggests that while CD300f deficiency may lead to reduced microglial reactivity to and engulfment of dying/dead cells, this phenotype is unlikely confounded by alterations in peripherally derived soluble factors. Such results underscore the regulatory role of CD300f in microglial phagocytosis.

To examine the potential role of CD300f in sensing tissue injury cues by other cell types, such as resident meningeal myeloid cells, we analyzed the production of chemokines and neutrophil and macrophage recruitment to the meninges following mTBI. Neutrophils express CD300f and typically serve as early responders to tissue injury in both the peripheral and CNS ^34^. Following mTBI, neutrophils participate in dead cell clearance and tissue repair within the meninges, with little or no effect to brain parenchymal damage ^28^. The absence of CD300f resulted in a reduction in the number of Ly6G+ neutrophils localized to the lesion site of the meninges at 5 hours post mTBI (Suppl. Fig. 1C-D). No differences in monocyte recruitment were observed (Suppl. Fig. 2E-F). This was confirmed by intravital two-photon microscopy of the meningeal space after mTBI (VIDEO 6) where we observed a substantial influx of Gr1+ cells to the WT meninges that was significantly reduced in CD300f^-/-^ mice at 5 hours after mTBI. Interestingly, this phenomenon was not accompanied by an increase in meningeal cell death (Suppl. Fig. 1G), suggesting that in our model, neutrophils do not participate in dead cell clearance/removal in the meninges, in contrast with previous reports ^28^. The diminished recruitment of neutrophils may stem from reduced detection of tissue damage by resident macrophages, leading to a decreased production of chemokines. We assessed the expression levels of neutrophil recruitment chemokines in brain and corresponding meninges 5 hours post mTBI in WT and CD300f^-/-^ brains, revealing a significant decrease in the mRNA of the neutrophil chemoattractant *cxcl1* and a trending reduction in *ccl3,* another neutrophil chemoattractant (Suppl. Fig. 2H). Furthermore, we investigated whether the reduction in neutrophil recruitment to the lesion site within the meninges would impact meningeal vascular repair. Interestingly we found no differences in revascularization rates in CD300f^-/-^ compared to WT meninges (Suppl. Fig. 2I-J). Collectively, these data suggest that the tissue-resident CD300f^-/-^ meningeal macrophages may inadequately sense tissue damage resulting in reduced production of neutrophil recruitment cues.

We subsequently opted to induce a more severe model of focal TBI, the controlled cortical injury (CCI), to evaluate the neurological impact of CD300f deficiency on responses to tissue damage and apoptotic cell sensing by microglia. Leveraging the parallel rod floor test, previously validated for the short- and long-term evaluation of motor coordination deficits post CCI ^35,36^, we followed lesioned mice for 90 days and observed impaired functional recovery in CD300f^-/-^ mice throughout the entire evaluation period compared to WT mice (Fig. 3A). There were no differences in spontaneous locomotor activity between the two groups (Suppl. Fig. 3A). Intriguingly, CD300f^-/-^ mice exhibited a trend towards a reduction in the lesion volume at 5 days post-lesion (Supp. 3B) and a significant reduction in the area and volume of the lesion at 90 days post-CCI (Fig 3B-D). Additionally, NeuN staining revealed a higher prevalence of neuronal bodies in the perilesion cortex of CD300f^-/-^ brains compared to WT mice at 90 days post-injury (Fig 3E, F). We then assessed the morphology of microglia in the perilesional cortex, as previously described ^37^, and observed a reduction in the ameboid/phagocytic microglia in CD300f^-/-^ brains (Fig. 3G, H). This finding aligns with the hypothesis that CD300f^-/-^ microglia fail to effectively sense DAMP cues and adopt a phagocytic morphology, thereby impeding the clearance of dead cells and potential debris following a sterile injury.

**Figure 3.**
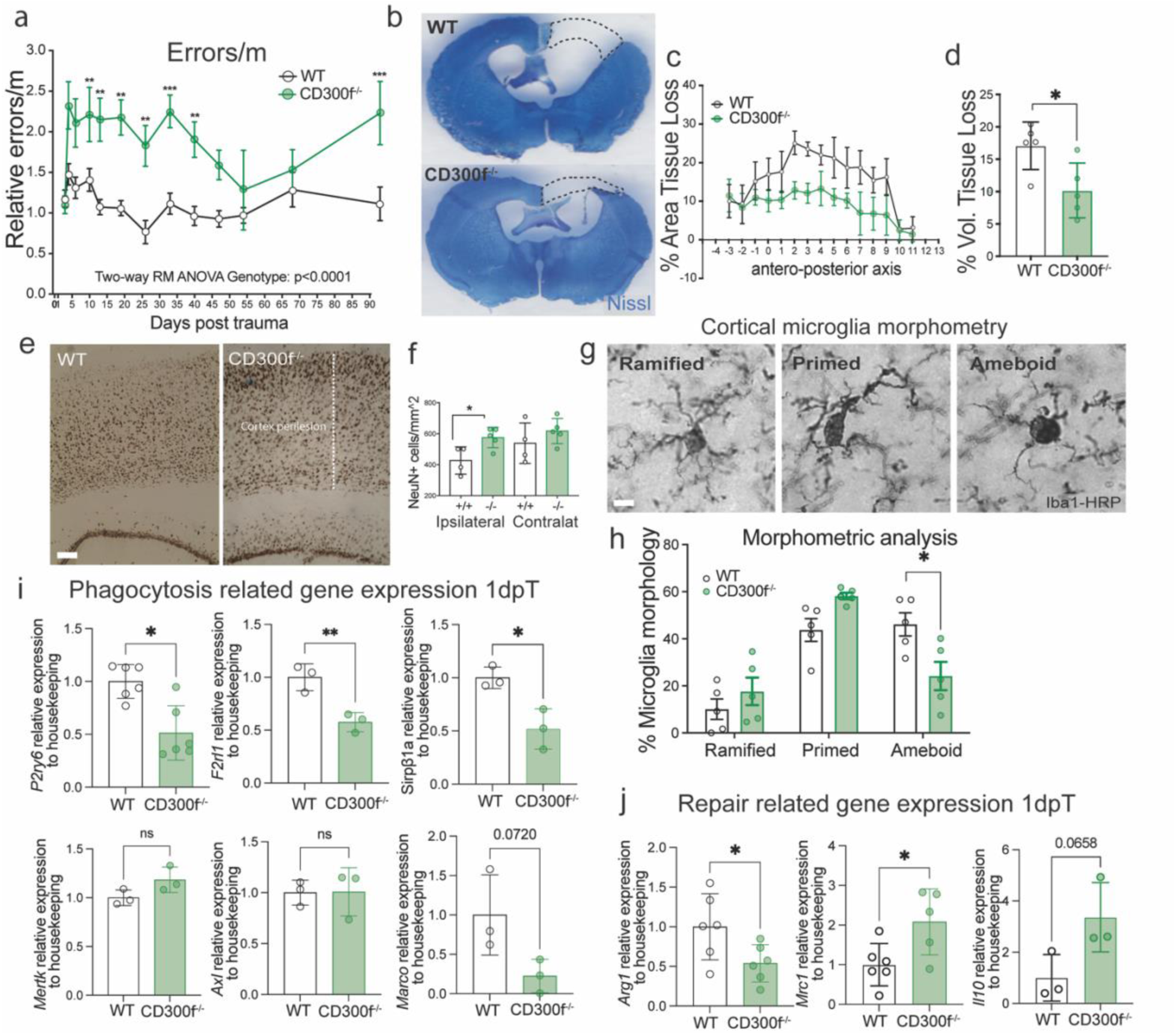
CD300f receptor impacts functional recovery and tissue loss/repair after severe TBI. **a.** Motor coordination impairment in male CD300f^-/-^ mice from 1-90 days after CCI, measured by the parallel rod floor test. The results were quantified as the number of errors committed per meter in a 10-minute period. Values are relative to the average of errors in each group at day 0 and normalized to the total distance travelled at day 0 before CCI. Two-way ANOVA, **p<0.01, ***p<0.001. n=13 animals per group. **b**, Nissl staining was performed to compare WT and CD300f^-/-^ brain tissue loss at 90dpl. The dotted line indicates the boundary between contralateral tissue volume and ipsilateral (remanent) tissue volume, representing tissue loss. **c**, Stereological analysis at 90 dpl demonstrates that CD300f^-/-^ animals have a reduction in the lesion area and **d**, volume calculated from remaining tissue compared with contralateral hemisphere. Two-tailed Student’s t-test, *p<0.05. n=5. **e**, Representative images and **f**, quantification of NeuN+ neuronal staining in the perilesioned cortex at 90 dpl show that CD300f^-/-^ animals have a higher number of surviving neurons compared to WT animals. Scale bar 100 μm. Two-tailed Student’s t-test *p<0.05. n=5. **g**, Representative images depict CD300f^-/-^ and WT Iba1^+^HRP microglia in the perilesioned cortex at 90 dpl, showing ramified, primed, and ameboid microglial morphologies. Scale bar 6 μm. **h**, Morphometric analysis show a reduction in ameboid microglia in CD300f^-/-^ brains at 90 dpl compared to WT. Data are expressed as the average percentage of microglial states per mm^2^. Two-way ANOVA, *p<0.05. n=5 animals per group. **i, j.** RT-qPCR analysis was conducted to examine the expression of **i**, phagocytic/scavenger related markers and **j**, repair related markers in CD300f^-/-^ and WT lesioned cortex at 90 dpl. Two-tailed Student’s t-test, *p<0.05 and **p<0.01. n=3-4 (WT) and n=3-6 (CD300f^-/-^).

Analysis of gene expression at 1 dpl (Fig. 3I) demonstrated a strong reduction in *P2ry6 mRNA* in CD300f^-/-^ brains, which suggests a diminished microglial phagocytic response to damage-associated alarmins, as demonstrated previously using the same mTBI model ^38^. In addition, *Sirpβ1a*, a phagocytic receptor in microglia/macrophages that contributes to the phagocytosis of dead cells and debris ^39^, and *F2rl1*, a receptor that promote chemotaxis, phagocytosis and oxidative burst in macrophages and microglia ^40^, were significantly reduced in CD300f^-/-^ brains. While no changes were observed in the mRNA of two key TAM receptors *Axl* and *Mertk,* involved in the phagocytosis of apoptotic cells by microglia, *Marco* showed a strong tendency to be downregulated. Additionally, we observed an increase in pro-regenerative/pro-fibrotic microglia/macrophages in the context of CD300f deficiency, as evidenced by the upregulation of *Mrc1* (CD206) and the trend in augmentation of *Il10* expression (Fig 3J). Although a reduction in *Arg1*, an anti-inflammatory marker on myeloid cells, was noted, no changes in classical pro-inflammatory cytokines such as *Il1β*, *Ifnγ*, *Tnfα*, or *Il6* were found in CD300f^-/-^ brains (Suppl. Fig. 3C), suggesting that CD300f deficiency does not induce an increase in generalized inflammation. Furthermore, while we observed a decrease in the expression of the neutrophil chemoattractant *Ccl3* at 1 day post CCI, there were no changes in *Cxcl1* (Supp. 3D).

Taken together, these data confirm that CD300f deficiency alters microglial phenotype, reducing long term microglial phagocytic programs and which results in enhanced functional compromise after a TBI.

## Discussion

Understanding the microglial sensome and the signals that coordinate microglial machinery towards a tissue injury, apoptotic cells or an infection is crucial, as these constitute critical triggers for the establishment of different cell states that will ultimately determine outcomes. The key finding of our work shows that CD300f lipid-binding immune receptor is essential for the in vivo microglial detection and directed polarization towards a tissue injury or apoptotic cells. Moreover, we show that this immune receptor is important for the in vivo phagocytosis of apoptotic cells and for the induction of adequate cellular processing machinery.

Two functionally and mechanistically distinct forms of microglial motility have been described. Directed motility to a laser-induced tissue lesion or a source of ATP is mediated by P2Y12 receptors ^41,42^. In fact, microglia from mice lacking P2Y12 receptors exhibit normal baseline motility but are unable to polarize, migrate or extend processes toward a specific stimulus ^41^. On the other hand, microglial ramification and surveillance of the brain are fundamentally dependent on the tonic activity of THIK-1 channels and do not require P2Y12 activity ^42^. The deficiency of CD300f did not alter the morphology or baseline surveillance of microglial cells in the brain, however it strongly impaired directed motility towards a tissue injury. The impaired extension of microglial processes towards a laser injury, with no affectation of the basal process motility, was also observed in TREM2 knockout mice ^43^, again suggesting functional convergence between these two receptors. One limitation of our study is that the exact niche or cells lesioned are not clear, which could explain the important variation in the lesions observed. Future studies should analyze if there are differences when for instance a blood vessel is injured, or if a neuron versus an astrocyte lesion induce the release of different DAMPs that elicits differential microglial responses. In relation to the mechanism underlying the impaired directed motility, no alteration in P2RY12 or the THIK-1 (*Kcnk13)* mRNA were described in vivo in CD300f deficient microglia ^12^, suggesting that CD300f may modulate directly the P2RY12 activation or indirectly some of its downstream effects to suppress directed motility. In fact, P2RY12 has been shown to be modulated by phosphorylation ^44^. In this line, CD300f has been shown to inhibit ATP-mediated mast cell activation probably by inhibiting P2Y7R by a mechanism that depends on its cytoplasmic inhibitory ITIM motifs that recruit phosphatases SHP1/SHP2 ^45^. Despite its inhibitory capacities, CD300f can also act as an activating receptor and is necessary for the full response of macrophages to IL4 ^46^. In this regard, CD300f could potentially be necessary for co-stimulating P2YR12 responses. Alternatively, CD300f may contribute to sensing DAMPs and PAMPs and coordinate the immunometabolic reprogramming necessary for orchestrating the many different functional changes required for the directed motility, secretion of cytokines, phagocytosis of debris and apoptotic cells, and accommodating the intracellular machinery for processing of phagocytosed materials. In accordance with this hypothesis, CD300f deficient microglia under inflammatory conditions have been shown to display strongly reduced metabolic fitness, not only with reduced basal glycolysis and respiration, but also being unable to reprogram its metabolism towards the induction of compensatory glycolysis during mitochondrial inhibition ^12^. Moreover, the same study demonstrated reduced expression of many autophagy related genes and of many lipid metabolism genes such as *Abcd1*, *Abcg1* and many β-oxidation enzymes and the carnitine shuttle transporter *Cpt1*. This is in accordance with the reduced capacity for processing apoptotic cells observed here in CD300f^-/-^ microglial cells in vivo after a mTBI or after injection of apoptotic thymocytes, or in vitro by CD300f deficient BMDM.

Apoptotic cells release specific metabolites, while retaining their membrane integrity ^47^. In this way, the apoptotic metabolite secretome is a regulated process and not the result of simply a passive emptying of cellular contents ^47^. Microglia deficient in CD300f failed to detect apoptotic cells in vivo and, in addition, showed a reduced capacity for phagocytosing apoptotic cells in vivo and in vitro. Some evidence suggests that CD300f is important for the phagocytosis of apoptotic cells, at least in vitro ^18^. The microglial P2Y_6_ receptor (P2RY6) – activated by extracellular UDP released by stressed neurons – is upregulated in microglia during in vivo neuronal death ^31^ and is required for microglial phagocytosis of neurons ^31,48^. Loss of P2RY6 strongly attenuated spontaneous microglial calcium signaling and was associated with reduced pro-inflammatory cytokine production, deficits in microglial phagocytosis, and enhanced CA3 neuron survival after epileptogenesis ^30^. It has also been shown that the inhibition of microglial P2RY6 prevented the transition of ramified microglia towards phagocytic microglia and increased the number of dead cells in the brain parenchyma in the same mTBI model used here ^38^. We observed a strong tendency towards downregulated mRNA for P2RY6 in naïve microglia in the brain of CD300f-/-mice, an effect that was more pronounced after 1 day after a CCI. Thus, the lack of P2RY6 in CD300f^-/-^ microglia could be an important part of the mechanism that includes the decreased UDP-dependent sensing of apoptotic cells and the reduced phagocytosis of injured neurons both after mTBI and after a CCI, and the increased number of surviving neurons at 3 months after the CCI. Moreover, it was reported that CD300f^-/-^ microglia express lower levels of the phagocytosis mediator *Spp1* mRNA in the naïve brain, and we show here that 1 day after the CCI there is also reduced expression of other phagocytosis related genes such as *Sirpb1a* and *Marco*, which could further contribute to decreased phagocytotic capacity. Finally, activation of CD300f by PS on apoptotic cells leads to its association to PI3K and to the downstream activation of Rac/Cdc42 GTPase, that mediates changes of F-actin that drive apoptotic cell efferocytosis ^18^. Thus, in the absence of CD300f, the malfunction of this mechanism would also contribute to the decreased phagocytosis.

Following the primary injury that occurs after a TBI, extensive and long-lasting tissue damage and functional compromise is prolonged through a complex cascade of events referred to as secondary injury. For many years, neuroinflammation has been proposed as an important process of the progressing secondary injury ^49^. However, neuroinflammation is an ill-defined process that can contribute to or mediate detrimental or beneficial effects. It is necessary to better understand the timing and complexity of the immune responses and cell states involved in TBI, complementing the transcriptomic point of view with in vivo experiments that could provide information at the functional level ^50^. TBI produces elevated levels of DAMPs and, if infection occurs during the days after the lesion, of associated PAMPs. These complex microenvironments will impact immune cells and determine their cell states. Immune receptors are critical regulators of the function of immune cells, particularly those that associates with adaptors with ITAMs such as DAP12/TYROBP and those with ITIMs in their cytoplasmic tail. Most of their ligands are transmembrane proteins, but an important subgroup are the lipid-binding immune receptors of the TREM ^51^ and CD300 families ^52^. Their ligands are not well characterized, in part due to its lipid nature, their diversity and their apparently unselective nature. This characteristic makes them putative DAMP and PAMP receptors which, in the context of the nervous system, may sense not only apoptotic cells and general cellular debris but also myelin debris ^53^. We report here that after a mild TBI, CD300f is important for sensing tissue damage and apoptotic cells and, in its absence, microglia display a strongly reduced capacity for extending processes towards the lesion and acquire a phagocytic amoeboid morphology. This progresses during the first 24 hours to an accumulation of dead cells both in the parenchyma and inside the few phagocytic microglia observed. To evaluate long-term neurological functional consequences of the TBI, we used a clinically relevant model of contusion injury, and observed that the absence of CD300f induced increased neurological deficits both at acute and sub-acute phases and at a long-term 3-month follow-up. Surprisingly, the total spared brain tissue of the lesioned hemisphere was increased in the CD300f^-/-^ mice. In addition, the number of neurons in the ipsilateral perilesional cortex was also increased when compared to control WT lesioned mice. This is in accordance with the reduced capacity of CD300f^-/-^ microglia and macrophages to detect and phagocytose stressed cells. Interestingly, we observed a reduced level of P2RY6 mRNA, and the absence of this receptor has been reported to inhibit microglial phagocytosis of stressed cells and result in increased neuronal numbers in models of epileptogenesis ^30^, after Aβ injection ^48^ or in a mouse TAU AD model ^48^. However, in opposition to the increased motor deficits we observed after the CCI, all these cases were associated with preserved neurological function. This suggests that CD300f has a broader regulation of microglial states that influence the outcome of TBI. For instance, CD300f has been shown in vitro to modulate synapse pruning due to its phosphatidylserine binding capacity ^15^. This could alter the neuroplasticity associated to neurorestoration after TBI and lead to increased motor deficits in the absence of increased tissue loss.

An important mechanism triggered by DAMPs is the activation of the inflammasome complex. Activation of pattern recognition receptors such as the NLR (NOD-like receptor) family or absent in melanoma (AIM) leads to autoactivation of caspase-1 and processing of pro-IL-1β and pro-IL-18 to their active forms. In patients, NLRP1, ASC and caspase-1 levels are increased in the CSF after severe TBI and are correlated with more unfavorable neurological outcomes ^54^. Recent data suggest a close relationship between NLRP3 inflammasome and CD300f. For instance, stimulation of BMDM with the PAMP LPS induced a stronger NLRP3 activation in the absence of CD300f ^13^. Moreover, NLRP3 knockout is neuroprotective in both TAU ^55^ and Aβ ^56^ Alzheimer’s disease models, and transcriptomic analysis showed that in both cases, among the very few differentially over-expressed genes was CD300f, suggesting that it may be an important neuroprotective receptor that is inversely regulated with NLRP3. Interestingly, the proposed mechanism involves a novel function of NLRP3 related to immunometabolism and not to IL1β production. The neuroprotection induced by NLRP3 knockout involves the upregulation of *Slc1a3* glutamate transporter that provides increased TCA cycle substrates to the mitochondria and in this way stimulates microglial metabolism ^56^. Interestingly, CD300f is upregulated in this condition, and CD300f has been shown to be important for proper microglial and macrophage immunometabolic fitness under inflammatory conditions ^13^. The unchanged levels of many pro-inflammatory factors, including IL1β, after the CCI further points towards other neuroprotective mechanisms of CD300f not related to classical inflammatory mechanisms, such as immunometabolic regulation of microglia.

Taken together, our data provide evidence for understanding the role of microglia and the function of CD300f immune receptor both in the naïve CNS and after an acute lesion. These data also contribute to understanding the broader mechanisms underlying pathologies in which this immune receptor has been involved, such as neuropsychiatric conditions ^12,17^, autoimmunity ^18,57^, psoriasis ^58^, allergies^59^, aging ^13^, multiple sclerosis ^22–24^ and TBI. Clearance of apoptotic cells has been linked to resolution of inflammation, but apoptotic cells are also generated under physiological conditions. In fact, a recent report shows that the identity of apoptotic cells induces different states in efferocytic macrophages in the context of IL4 ^60^, highlighting the importance of describing the type of receptors engaged by different apoptotic cells during this process. Thus, from a broader perspective, these data contribute to the comprehension of microglia and macrophage responses to local tissue signals that regulate the spectrum of functional states that they acquire within a tissue.

## Methods

### Rodents

C57BL/6J (B6), B6.129P-Cx3cr1tm1Litt/J (CX3CR1^gfp/gfp^) mice were obtained from The Jackson Laboratories. CX3CR1^gfp/+^ mice were generated by crossing B6 mice with CX3CR1^gfp/gfp^ mice in a closed breeding facility at the National Institutes of Health (NIH). B6.Cg-Cd300lftm1Jco/J (CD300f ^-/-^) mice were obtained from NIAID/NIH ^18^ and maintained at the NINDS facility under controlled environment. All mice were between 12–32 weeks of age. All rodents were housed under specific pathogen-free conditions and treated in accordance with Institutional Animal Care and Use Committee at the NIH. For Controlled Cortical Injury (CCI) experiments, adult (4- to 5-month-old) C57BL/6 (Charles River) WT mice and CD300f^−/−^ (Genentech ^61^) mice were crossed, regenerated and then bred for experiments at the SPF animal facility of Institut Pasteur de Montevideo-Uruguay. Animals were maintained under controlled environment (20 ± 1 °C, 12-h light/dark cycle, free access to food and water).

### Skull Thinning & Compression Injury

mTBI injuries were performed as described previously ^62^. Mice were anesthetized by i.p. injection of ketamine (85 mg/kg; Ketaset), xylazine (13 mg/kg; AnaSed), and acepromazine (2 mg/kg; VET one) in PBS, and body temperatures were held at 37°C. An incision was made to expose the skull bone, and a metal bracket was secured on the skull bone over the barrel cortex (2.5 mm from bregma × 2.5 mm from sagittal suture). A 1 × 1 mm cranial window was quickly thinned over 1– 2 minutes using a dental drill with a 0.5 mm burr to a thickness of ∼20–30 μm. Once thinned, a microsurgical blade was used to lightly compress the skull bone.

### Intravital two-photon laser scanning microscopy and laser-induced injury

CX3CR1^gfp/+^ x CD300f ^-/-^ or ^+/+^ male and female mice were used to visualize myeloid cells through a thinned-skull window in naive mice and following mTBI. Compressed and control mice were imaged using a Leica SP8 2-photon microscope equipped with an 8000-Hz resonant scanner, a 25× collar-corrected water-dipping objective (1.0 NA), a quad HyD external detector array, and a Mai Tai HP DeepSee Laser (Spectra-Physics) tuned to 905 nm. For the demonstration of dead cells, a solution of 1 mg/ml propidium iodide (ThermoFisher P3566) was used for transcranial incubation and intraperitoneal injection. For blood vessel visualization, 100 μL of 1 mg/mL Evans blue (MilliporeSigma) dissolved in PBS was injected i.v. immediately following surgical preparation. For in vivo neutrophil recruitment analysis, mice received an i.v. injection of 488-anti-GR1 (BD 108406) 5 hours after mTBI immediately prior to imaging. For all imaging, artificial cerebral spinal fluid (aCSF) (Harvard Apparatus, 597316) was used to submerge the lens above the thinned skull.

Laser injuries were performed applying 100% of Mai Tai laser tuned at 1050nm in a ROI of 17 μm for 2 minutes and imaged using Insight laser tuned at 905 nm. Three-dimensional time-lapse movies were captured as Z stacks of 1.0 µm for laser injuries, or 2 μm for phagocytosis assays, to a depth of 20 μm and 70 μm respectively. Compressed and laser injury time lapse videos were acquired with 1 min intervals, and 22 seconds intervals between 3D stacks respectively. For the quantification of microglial response to cerebral laser ablation, a region of interest (ROI) around the injury site was defined using IMARIS software. The volume of adjacent microglial processes intersecting the ROI was measured over a 30-minute period. The laser ablation experiments were conducted using 6 WT and 6 CD300f^-/-^ male and female animals, including a total of 10 and 12 lesions respectively. Lesions were performed in at least one WT and one CD300f^-/-^ mouse per day in five different days. Two lesions from one WT animal were not used as microglial processes retracted instead of reacting towards the lesion. One lesion from a CD300f^-/-^ animal was not used as it produced a massive microglial reaction not seen in other lesions, and thus was not considered. It must be noted that, despite careful methodological design and execution, there are important differences in the lesions observed within each group, which suggests that additional sources of variations exist.

### Glial limitans leakage assay

To assess the integrity of the glia limitans, a compression injury was induced in CX3CR1^gfp/+^ x CD300f ^-/-^, ^+/+^ male and female mice. After 5 hrs, 200 μL sulforhodamine 101 (SR101) (1 mM, S7635, Sigma-Aldrich) dissolved in PBS was applied to the skull for 12 minutes. SR101 was then washed several times with aCSF over a 5-minute period. A 3-dimensional 300 μm Z stack (1 μm step size) was then captured using the 2-photon microscope. Z stacks were started just above the skull bone. For quantification, Z stacks were cropped to 250 μm to remove the skull bone. The mean fluorescence intensity (MFI) of the channel corresponding to SR101 signal (562–650 nm) was then calculated using Imaris version 9.6 image analysis software.

### CCI

Adult male WT and CD300f^−/−^ mice (20 to 25 g) were injured using a controlled cortical impactor (CCI, PinPointTM Precision Cortical Impactor Model PCI3000 Hatteras Instruments, Cary, NC, USA) as described in ^36^. Briefly, mice were anesthetized using isofluorane chamber and mounted in the prone position in the stereotaxic frame of the cortical impactor device. A midline incision was performed, and using a dental drill, a 3-mm round craniotomy hole was created on the right hemisphere between bregma and lambda, immediately adjacent to the coronal suture, and 0.5 mm proximal to the sagittal suture. The resulting 3-mm diameter bone flap was removed. The center of the CCI tip was placed over the right primary motor and somatosensory cortex (coordinates for the tip center in mice, L: 1.75 mm, AP: −0.75 mm) and the CCI was performed with a velocity of 2 m/sec, a contusion dwell time of 150 msec, and a depth of 2mm.

### Histology and Immunohistochemistry

Mice were perfused with a solution of 4% paraformaldehyde (PFA) and placed in 4% PFA for 2 hours, washed and placed in 25% sucrose solution for 2 days for cryopreservation. Brains were then embedded in OCT medium and frozen over dry ice, and 30-100 μm coronal sections were made using a cryostat Leica CM1850 UV. Brain sections were blocked and stained on slides as described below.

### Tissue preparation

To examine the number of dead cells in the brain parenchyma after compression injury, the dead cells were labeled by transcranial propidium iodide (PI, 1mg/ml Thermo Fisher P3566) application for 45 minutes in aCSF. This was followed by a single wash with aCSF and imaging. The mice were perfused with 4% PFA, and the brain tissue processed as described below.

For immunostaining of brain slices and whole-mount meninges, the tissue was immersed in a block-permeabilization solution containing 10% FBS, 0.5% Triton-X-100 in PBS for 1 h at room temperature. Subsequently, the tissue was stained for antigens with anti-mouse antibodies in 1% FBS and 0.5% Triton-X-100 in PBS at RT overnight. For brain slices, rabbit anti-Iba1 (1:700, 019-19741, Wako Chemicals) was incubated ON at RT and detected with goat anti–rabbit Alexa Fluor 488 (1:1000, Thermo Fisher, A-11034). Brain sections were incubated with 1:500 DAPI (1 mg/mL, D9542, Sigma-Aldrich) in PBS for 10 minutes at room temperature before washing and mounting on charged glass slides.

### Dead cell detection and quantification

A single section with the highest density of propidium iodide staining was imaged through entire Z plane using an Olympus FV1200 laser-scanning confocal. DAPI+ “surfaces” were created, and dead cells were enumerated as DAPI+ cells that colocalized with propidium iodide. Dead cells were normalized for each experiment by averaging the injury in the WT group and calculating the fold change for each individual sample relative to the average. Data is expressed as a fold change of CD300f+/+.

Immunostaining of brain and mouse whole-mount meninges was performed as previously described ^63^ with modifications. Skull caps, with intact dura mater, were dissected and placed in 2% paraformaldehyde for 16–18 h at 4 °C. The fixed tissue was then washed with PBS three times in PBS for 5 min per wash. Under a dissecting microscope, the dura mater was scored carefully with forceps from the skull cap, leaving the meninges intact, in a Petri dish containing PBS. Meninges were then stored at 4 °C in PBS in a 24-well plate until they were used for immunostaining.

### Meningeal whole mounts staining

A combination of the following primary antibodies was used for immunostaining of whole-mount meninges: anti-Ly6G (1:200 BioLegend 127626), anti-CD11b (1:200 BioLegend 101220) anti-Laminin (1:200 abcam, ab11575). Following staining, tissue was washed three times in PBS for 5 min per wash at room temperature. Using a paintbrush, the dura mater was flattened on a glass slide and mounted with FluorSave Reagent (Millipore Sigma). Slides were left in the dark at 4 °C for 1 h before imaging. Laminin-lectin paradigm of vascular repair 7 days after TBI was performed as described previously on ^64^

### Confocal Microscopy

Tile scan imaging of brain slices or whole-mount meninges was carried out on an Olympus FV1200 laser-scanning confocal microscope equipped with four detectors, six laser lines (405, 458, 488, 515, 559 and 635 nm) and five objectives (4×/0.16 NA, 10×/0.4 NA, 20×/0.75 NA and 40×/0.95 NA, and chromatic aberration-corrected 60×/1.4 NA). Images were analyzed using Imaris v.9.3 software (Bitplane).

### Histology – Nissl staining

Parallel coronal cryostat sections (30µm thick) of the entire brain, separated by 300µm (every 11th section) were stained for Nissl as described in ^25^ and used for the quantification of the lesioned area and the total area of the hemisphere. Quantification of the lesioned Nissl pale area was performed using the Fiji (National Institutes of Health, USA, http://rsbweb.nih. gov/ij/index.html) with parallel microscope observation (using 40× magnification) by two independent researchers blinded for the treatment administration as described previously. The slice 0 was established as the coordinate in the in the cephalon-caudal axis where lateral ventricles are visualized. The remanent area of the lesioned hemisphere and the total contralateral hemisphere area were used to calculate the lesion conserved tissue area and the inverse value (area of tissue loss). The data was expressed as “% of tissue loss” to correct for the possible edema effect.

Immunohistochemistry was performed on coronal sections to assess the number of neurons and morphological activation states of microglia in the perilesional cortex, as described previously ^65^. Briefly, the endogenous peroxidase was inhibited with 10% methanol in 30% PBS and 2% H_2_O_2_ for 10 minutes, the tissue background was blocked for 1h, and sections were washed with 1x PBS and incubated overnight at room temperature with rabbit anti-Iba-1 (1:700; Wako Chemicals) or anti-NeuN (ABN78, Millipore). Then, sections were washed in 1× PBS-Triton x100 1% three times and incubated with HRP-anti-rabbit IgG antibody for 2 h at room temperature and incubated with 3,3′-diaminobenzidine (DAB) 0.5mg/mL in PBS + H_2_O_2_ for colour development. Fiji software was used to count and classify the number of cortical microglia and neurons, and for microglial morphologic phenotypes (ramified primed and hypertrophic/ameboid). The analyses were performed by a blind investigator.

Microglial phenotypic classification was based on the length and thickness of the projections, the number of branches, and the size of the cell body, as previously described ^66^. Briefly, ramified microglia had long thin processes (>650 μm in length), a small cell body volume (10 μm3), and many branches (from 20 to 30). Primed microglia had medium-length processes (300–550 μm in length), larger cell body volumes (50–75 μm3), and from 20 to 30 branches. Ameboid microglia had short thick processes (∼200 μm in length), a larger cell body volume (80–100 μm3), and very few branches (∼10). Ramified microglia had a nonactivated (resting) morphology, whereas primed and ameboid microglia had an activated and ameboid-like morphology. Data was expressed as a % of all microglia visualized in each slice. An average of 5 slices was use per CD300f-/- and +/+ lesioned animal.

### Flow Cytometry

mTBI was performed 24 hours prior to the phagocytosis assay on CD300f^-/-^ or ^+/+^ mice under aseptic conditions. Three to five brain punches of 2 mm² per group were pooled together, and microglial cells were then dissociated into a single-cell suspension using the Miltenyi Tumor Dissociation Kit (130-096-730), GentleMACS 25C Tubes (130-093-237), and GentleMACS Octo Dissociator. The cells were isolated using anti-human CD11b beads (Miltenyi 130-093-636) and myelin removal beads (Miltenyi 130-096-733). All directly conjugated antibodies purchased from BioLegend. Single-cell suspensions were were acquired using a BD FACSymphony A5 flow cytometer and analyzed using FlowJo software version 10.9.0.

### Apoptotic thymocytes assays

Thymus dissection and single cell suspension were performed by mashing the tissue through a 70 μm pore cell restrainer, followed by ON treatment with dexamethasone 0.1 μM to induce apoptosis. Apoptotic thymocytes were then labeled with CMTPX dye (Thermo Fisher C34552) as indicated by manufacturer and injected intracerebrally using a stereotactic device at a concentration of 50,000 cells in 2 μl. After a 3-hour incubation period, analyses were conducted using either flow cytometry FACSymphony A5 or intravital two-photon microscopy as described above.

For bone marrow derived macrophages (BMDM) phagocytosis experiments, cells were obtained as described on^13^. Briefly, cells were isolated from adult female and male WT and CD300f^−/−^, bone marrow was flushed from the femur and tibia with DMEM/F12 (Gibco, Cat # 10565018) supplemented with Penicillin/Streptomycin and 10% FBS (complete DMEM/F12). Cells were plated on to 100 mm bacteriological Petri dish with complete DMEM/F12 and 20 ng/mL recombinant mouse macrophage colony stimulating factor protein (M-CSF) (Biolegend, Cat #576406). On day 4, cells were subcultured in XFe24 plates at 100,000 cells per well or at 200.000 cells per well in XFe96 plates. Apoptotic thymocytes were stained with CFSE (Thermo Fisher C34554) and added to BMDM cultures for 15’ or ON. Cells were collected for flow cytometry acquisition and analysis on FlowJo as previously described. Phagocytic BMDM were determined by analyzing the cells that uptake CSFE+ apoptotic thymocytes.

### Gene expression analysis

CCI mice at 1-day post-lesion and mTBI at 5h post-lesion were perfused with cold PBS, followed by brain dissection and snap freezing. Superficial cortical tissue and meninges were removed from a small area surrounding the TBI lesion and placed in RPMI. Tissue was mechanically homogenized using sterile zirconia/silica beads (Biospec). Homogenized tissue was spun down at 12,000 g at 4 °C and resuspended in Trizol (Thermo Fisher). RNA extraction was performed using RNeasy Plus Micro Kit (50) (cat# 74034), followed by reverse transcription (RT) and cDNA synthesis. Using a CFX96 Real-Time PCR machine (BioRad Laboratories), real-time PCR was performed with 40 ng of DNA per sample using SYBR Green (Applied Biosystems) or water (non-template negative control). For CCI, pre-made PrimePCR™ plates (Bio-Rad) targeting inflammation and phagocytosis pathways were utilized according to the manufacturer’s instructions. Briefly, 10–20 ng of cDNA and SYBR green reagent (Bio-Rad) were added to lyophilized primers in each well and the plate was read on a Bio-Rad CFX96 Real-Time with C1000 Thermal Cycler system. On each PrimePCR plate there were control wells for RNA quality and gDNA contamination. qPCR data was analyzed using Bio-Rad Gene Study software, with expression values relative to those of the control genes Gapdh and Tbp.

### Behavior Analysis

The short- and long-term effects of TBI were evaluated using the Parallel Rod Floor test (Stoelting) and the ANY-maze software, as previously described ^36,67^. The number of errors and distance travelled were automatically measured by the software, and errors per meter were normalized using the average of WT mice at baseline (day 0 pre-CCI). The results are expressed as fold change. Behavioral experiments were conducted during the illuminated phase of the cycle under dimly lit conditions using a red lamp and low noise. Behavior was monitored and videotaped by a blinded, trained observer, and the results were obtained accordingly.

### Statistical analysis

Continuous variables with normal distribution were evaluated using Student’s two-tailed t test, one-way analysis of variance (ANOVA) followed by Tukey’s post-hoc analysis, or two-way ANOVA for repeated measures followed by Sidak’s post hoc test, as appropriate. All statistical analyses and significance (P ≤ 0.05) were performed using GraphPad Prism 10.0.

## Supplementary figure legends

**Supplementary 1. No changes in microglial morphology or basal surveillance dynamics. a,** Confocal images of CX3CR1^+/GFP^/CD300f^-/-^vs WT CX3CR1^+/GFP^ naïve control brains showing microglia under homeostasis. No differences in morphological/functional parameters, such as sphericity (b) and volume (c) were detected. n=3 animals per group, 5 images per mice. Scale bar 20 μm. Two-tailed Student’s t-test.

**Supplementary 2. CD300f deficiency affects meningeal neutrophil recruitment. a, b.** No difference in glial limitans leakage was observed after mTBI in CD300f-deficient mice compared to WT, as indicated by SR101 assay. Data representative from 3 independent experiments. n= 5 (WT) n=4 (CD300f^-/-^). Scale bar 30 μm **c-g.** Meningeal whole mounts show a decrease in neutrophil infiltration (**c, d**) into the site of the lesion in CD300f^-/-^ mice 5 hours after mTBI. Data pooled from 6 independent experiments. n=15 (WT) n=14 (CD300f^-/-^). Scale bar 1000 μm and 300 μm. Ly6G (green), lectin (red). No differences monocyte counts (**e**) or in meningeal cell death (**g)** were observed, CD11b (white) n=6. PI (red) n=11 (WT) n=10 (CD300f^-/-^). Scale bar 200 μm. Two-tailed Student’s t-test, ****p<0.0001. **h.** Reduction in chemokine *Cxcl1* expression levels in brain punctures analyzed 5h after mTBI. No differences in expression levels of *Ccl*3 or *Mrc1.* n=3-4 (WT) n=4 (CD300f^-/-^). Two-tailed Student’s t-test*p<0.05. **i, j,** Lectin-laminin assay performed seven days post-mTBI (red dotted line) show no differences in meningeal vascular repair between CD300f^-/-^ and WT mice, laminin (red), lectin (white). Scale bar 200 μm. n=4 mice per group. Two-tailed Student’s t-test.

**Supplementary 3. CD300f deficiency does not impact spontaneous locomotor activity or general pro-inflammatory gene expression. a.** Spontaneous locomotor activity measured by the distance traveled in the arena over a 10-minute period in CD300f^-/-^ and WT mice. Two-way ANOVA *p<0.05. n=13 per group. **b.** Lesion volume at 5 dpl in CD300f deficient mice. Two-tailed Student’s t-test. n=3 (WT) n=4 (CD300f^-/-^) **c.** Expression levels of acute inflammation and **d**, recruitment genes in CD300f^-/-^ mice at 1 dpl (+ naïve and 90 dpl in the case of *Ccl3*). Two-tailed Student’s t-test. n=3-12 (WT) n=4-12 (CD300f^-/-^) **e.** Heat maps of acute inflammation and phagocytic pathways obtained by qPCR premade plates. The asterisk in red show the DE genes. Two-tailed Student’s t-test. n=3 animals per group.

## Author contributions and acknowledgments

H.P. and M.L.N. conceived the study. M.L.N. performed most of the experiments. F.E., F.A.C., H.D.M Z.F and D.A. performed experiments. M.L.N. F.E. Z.F., and H.P. analyzed and discussed results. H.P. drafted the original manuscript and M.L.N. and Z.F. contributed to the edition of the final version. All authors have read and agreed to the published version of the manuscript. We thank the staff of the URBE, Facultad de Medicina UDELAR (Uruguay), and Laboratory Animals Biotechnology Unit (UBAL), Advanced Bioimaging Unit (UBA) and Cell Biology Unit of the Institut Pasteur de Montevideo (Uruguay).

