## Supplementary figures for "CD300f immune receptor is a microglial tissue damage sensor and regulates efferocytosis after brain damage"

**Supplementary 1. No changes in microglial morphology or basal surveillance dynamics.** Two-photon timelapse Scholl analysis for measuring microglial processes migration of individual animals is shown (a). Confocal images of CD300f$^{-/-}$CX3CR1$^{+/GFP}$ vs WT CX3CR1$^{+/GFP}$ naïve control brains showing microglia under homeostasis (b). No differences in morphological/functional parameters, such as sphericity (c) and volume (d) were detected. n=3 animals per group, 5 images per mice. Scale bar 20 µm. Two-tailed Student's t-test.

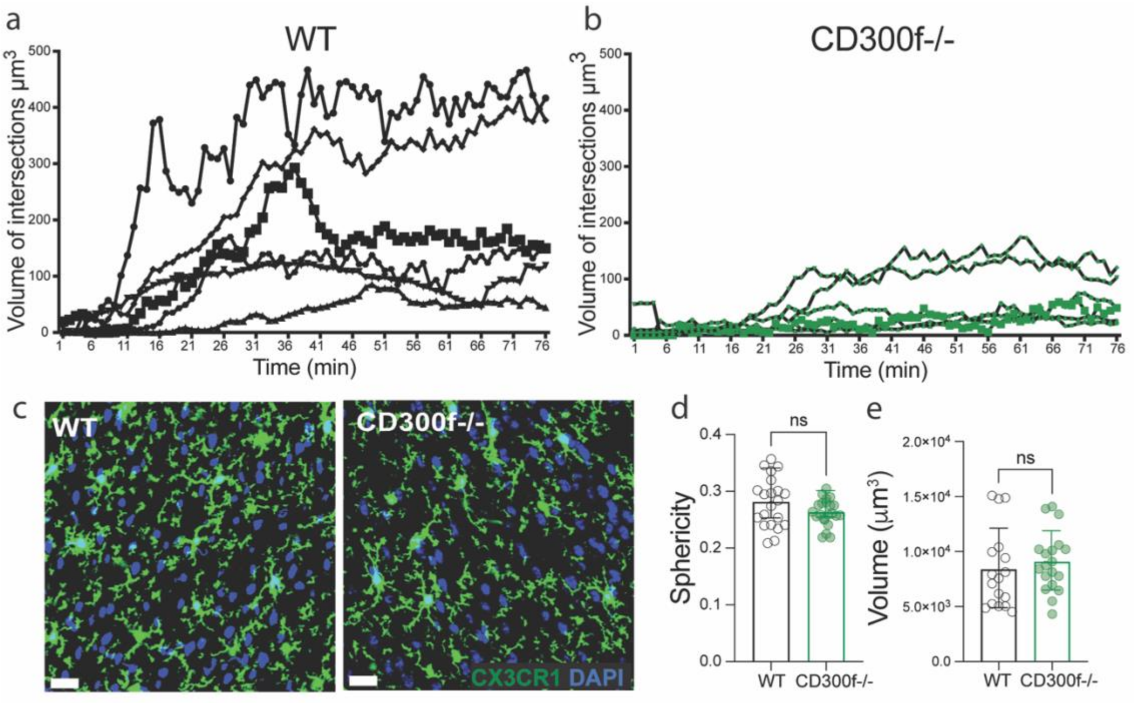

**Supplementary 2. CD300f deficiency affects meningeal neutrophil recruitment. a, b.** No difference in glial limitans leakage was observed after mTBI in CD300f-deficient mice compared to WT, as indicated by SR101 assay. Data representative from 3 independent experiments. n= 5 (WT) n=4 (CD300f$^{-/-}$). Scale bar 30 µm **c-g.** Meningeal whole mounts show a decrease in neutrophil infiltration (**c, d**) into the site of the lesion in CD300f$^{-/-}$ mice 5 hours after mTBI. Data pooled from 6 independent experiments. n=15 (WT) n=14 (CD300f$^{-/-}$). Scale bar 1000 µm and 300 µm. Ly6G (green), lectin (red). No differences monocyte counts (**e**) or in meningeal cell death (**g)** were observed, CD11b (white) n=6. PI (red) n=11 (WT) n=10 (CD300f$^{-/-}$). Scale bar 200 µm. Two-tailed Student's t-test, ****p<0.0001. **h.** Reduction in chemokine *Cxcl1* expression levels in brain punctures analyzed 5h after mTBI. No differences in expression levels of *Ccl*3 or *Mrc1.* n=3-4 (WT) n=4 (CD300f$^{-/-}$). Two-tailed Student's t-test*p<0.05. **i, j,** Lectin-laminin assay performed seven days post-mTBI (red dotted line) show no differences in meningeal vascular repair between CD300f$^{-/-}$ and WT mice, laminin (red), lectin (white). Scale bar 200 µm. n=4 mice per group. Two-tailed Student's t-test.

**Supplementary2:**

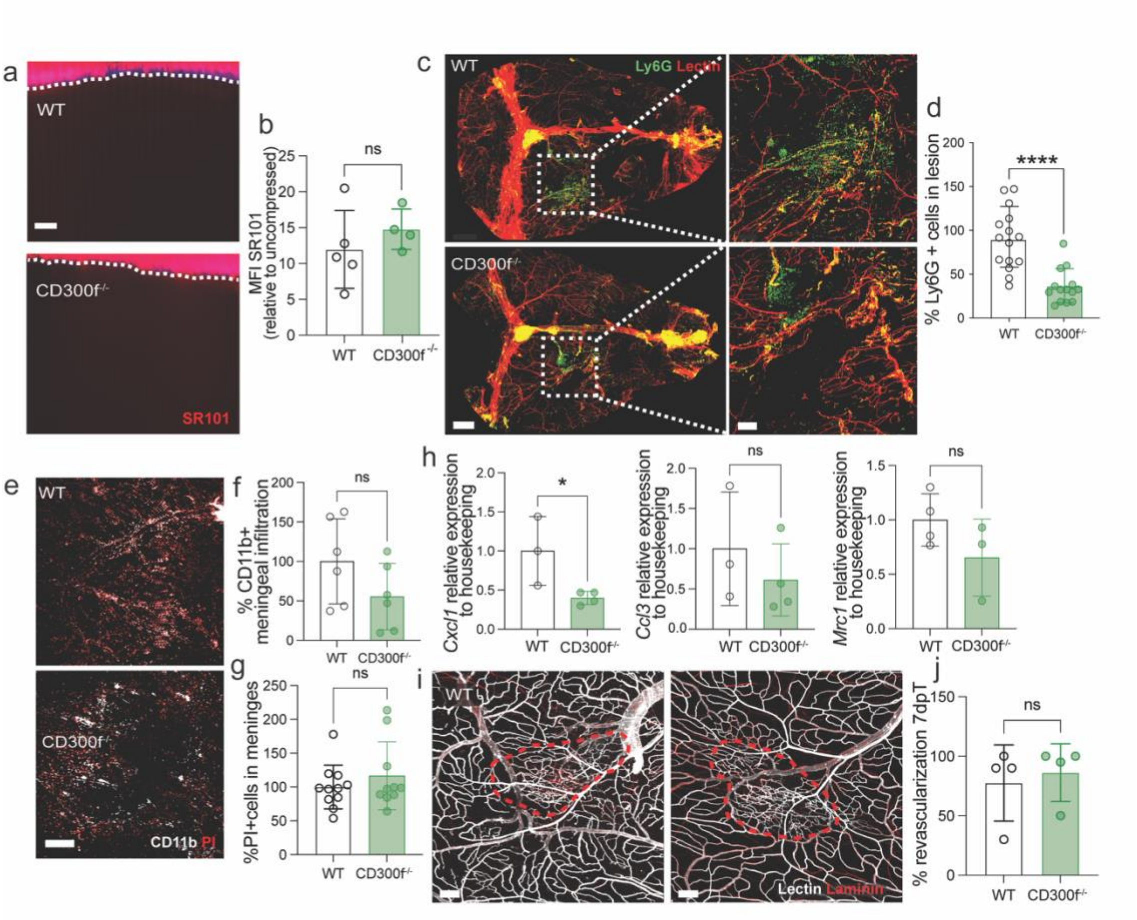

**Supplementary 3. CD300f deficiency doesn't impact spontaneous locomotor activity or general pro-inflammatory gene expression. a.** Spontaneous locomotor activity measured by the distance traveled in the arena over a 10-minute period in CD300f$^{-/-}$ and WT mice. Two-way ANOVA *p<0.05. n=13 per group. **B.** Lesion volume at 5 dpl in CD300f deficient mice. Two-tailed Student's t-test. n=3 (WT) n=4 (CD300f$^{-/-}$) **c.** Expression levels of acute inflammation and **d**, recruitment genes in CD300f$^{-/-}$ mice at 1 dpl (+ naïve and 90 dpl in the case of *Ccl3*). Two-tailed Student's t-test. n=3-12 (WT) n=4-12 (CD300f$^{-/-}$) **e.** Heat maps of acute inflammation and phagocytic pathways obtained by qPCR premade plates. The asterisk in red show the DE genes. Two-tailed Student's t-test. n=3 animals per group.

**Supplementary 3**

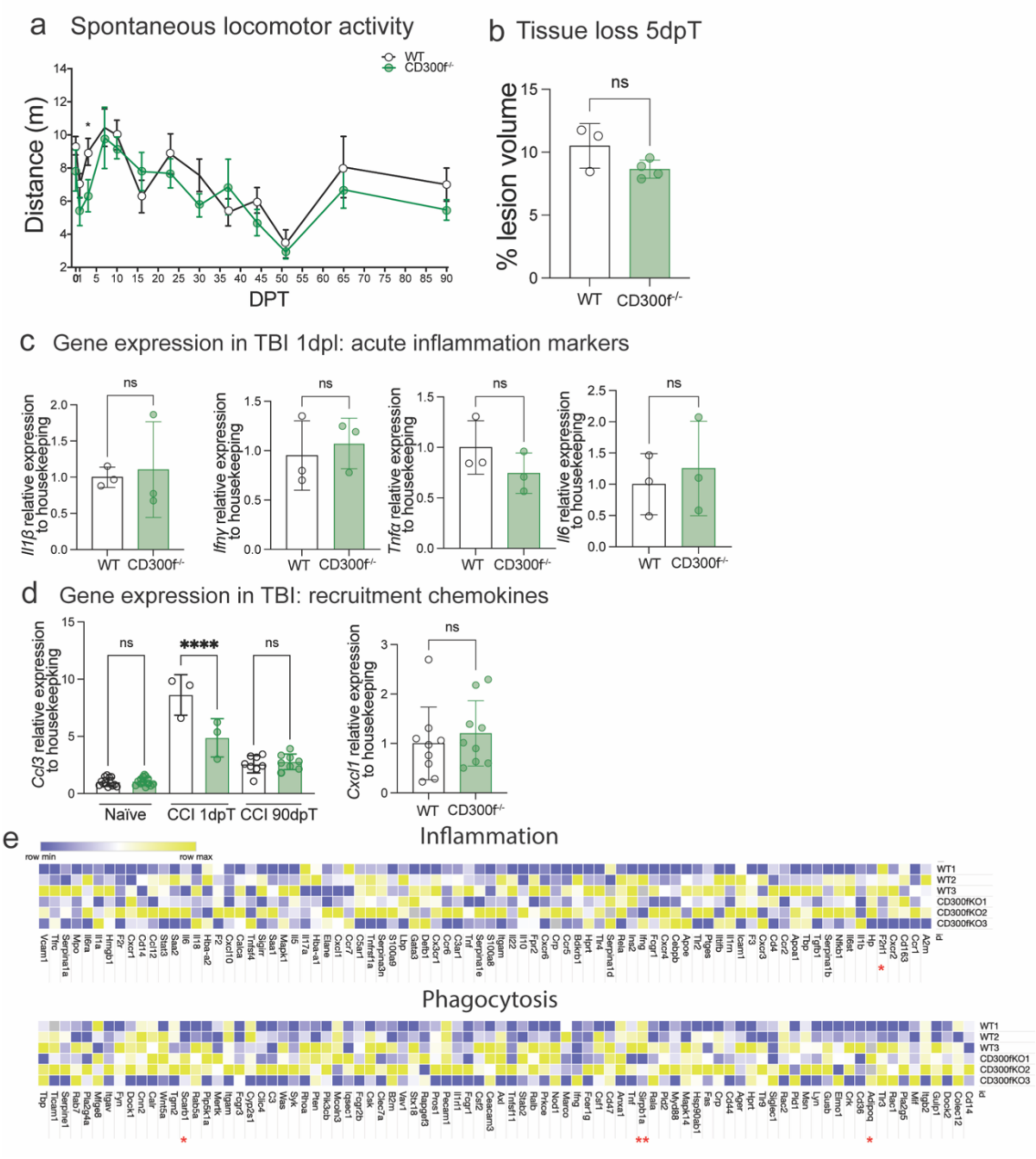
